# miTAR: a hybrid deep learning-based approach for predicting miRNA targets

**DOI:** 10.1101/2020.04.02.022608

**Authors:** Tongjun Gu, Xiwu Zhao, William Bradley Barbazuk, Ji-Hyun Lee

## Abstract

microRNAs (miRNAs) are a major type of small RNA that alter gene expression at the post-transcriptional or translational level. They have been shown to play important roles in a wide range of biological processes. Many computational methods have been developed to predict targets of miRNAs in order to understand miRNAs’ function. However, the majority of the methods depend on a set of pre-defined features that require considerable effort and resources to compute, and these methods often do not effectively on the prediction of miRNA targets. Therefore, we developed a novel hybrid deep learning-based approach that is capable to predict miRNA targets at a higher accuracy. Our approach integrates two deep learning methods: convolutional neural networks (CNNs) that excel in learning spatial features, and recurrent neural networks (RNNs) that discern sequential features. By combining CNNs and RNNs, our approach has the advantages of learning both the intrinsic spatial and sequential features of miRNA:target. The inputs for the approach are raw sequences of miRNA and gene sequences. Data from two latest miRNA target prediction studies were used in our study: the DeepMirTar dataset and the miRAW dataset. Two models were obtained by training on the two datasets separately. The models achieved a higher accuracy than the methods developed in the previous studies: 0.9787 vs. 0.9348 for the DeepMirTar dataset; 0.9649 vs. 0.935 for the miRAW dataset. We also calculated a series of model evaluation metrics including sensitivity, specificity, F-score and Brier Score. Our approach consistently outperformed the current methods. In addition, we compared our approach with earlier developed deep learning methods, resulting in an overall better performance. Lastly, a unified model for both datasets was developed with an accuracy higher than the current methods (0.9545). We named the unified model miTAR for miRNA target prediction. The source code and executable are available at https://github.com/tjgu/miTAR.

## Introduction

microRNAs (miRNAs) are a major type of small regulatory RNA that are ∼22 nucleotides (nts)(Bartel, 2004). They typically form complementary hybrid sequences with their targets, and act to repress gene expression at the translational level or cleave mRNAs at the post-transcriptional level(Bartel, 2004; Jonas and Izaurralde, 2015). It has been reported that miRNAs play key roles in a variety of biological processes and human diseases(Huang, et al., 2011), including cell differentiation and development, metabolism, proliferation and apoptosis, viral infection, tumorigenesis, diabetes, macro- or micro-vascular complications, neurological diseases. Thus, it is important to find the targets of miRNAs to better understand the function and regulation of miRNAs.

Advances in our understanding of the interactions between miRNAs and their targets has led to the development of many computational methods/tools to predict miRNA targets. The majority of these tools are based on common features of the miRNA:target interaction. Four features are widely used: sequence complement (especially in the seed region that is generally defined as a 6 or 7 nts sequence starting at the second or third nucleotide(nt) of the miRNA sequence), thermodynamic stability, target site accessibility and sequence conservation among species(Peterson, et al., 2014). Several widely used tools have been developed based on these features. For example, miRanda(Enright, et al., 2003) relies on sequence complementarity and binding energy; TargetScanS(Lewis, et al., 2005) relies on sequence complementarity in seed region; while PITA(Kertesz, et al., 2007) relies on target site accessibility. However, miRNA targets predicted by different methods and tools are inconsistent with one another. Furthermore, using known features limits the ability to predict novel or non-canonical miRNA targets, which have been determined to be prevalent(Agarwal, et al., 2015).

Recently, several deep learning methods were developed to handle unknown features and to improve the accuracy of prediction. MiRTDL(Cheng, et al., 2016) uses a convolutional neural network (CNN) to capture training sets features. Although CNNs can automatically assess feature importance, miRTDL is still based on known features. MiRTDL used 20 features from three categories: three conservative features, nine complementary features and eight accessibility features. Consequently, the information outside the 20 features cannot be captured. DeepTarget(Lee, et al., 2016) uses deep recurrent neural network (RNN) based autoencoders to learn sequence features for miRNAs and genes separately, and then uses a stacked RNN to learn the sequence-to-sequence interactions between miRNA and their targets. It supplies a way to predict the target of miRNAs without using any pre-defined features. However, RNN may not be efficient in learning spatial features that exist in miRNA:target interactions. In 2018 two newer deep learning methods were developed: DeepMirTar(Wen, et al., 2018) and miRAW(Pla, et al., 2018). DeepMirTar collects a set of 750 features and uses stacked denoising auto-encoders for miRNA target prediction. Although DeepMirTar significantly increases the number of features, it still depends on features derived from the four major feature types mentioned previously. MiRAW includes three major steps: two filter steps were used before and after a deep feed forward neural network step. The input for miRAW are the sequences of the concatenated miRNA and its target, which avoids pre-definition of the features. However, a feed forward neural network may not be efficient to capture the spatial and sequential features of the hybrid sequences of miRNA:target.

Encouraged by the higher accuracies and large training datasets collected from recent deep learning studies we developed a novel deep learning-based approach that integrates two major types of neural networks, CNNs and RNNs, to predict miRNA targets. CNNs are designed to learn spatial features; RNN are designed to learn sequential features (Zhang, et al., 2019; Zou, et al., 2019). By combining CNNs and RNNs, our approach has the advantages of learning both the intrinsic spatial and sequential features of miRNA:target. The inputs for our models are the primary sequences of miRNAs and genes, which can be easily obtained. We collected two datasets from the studies of DeepMirTar and miRAW as our training datasets. By training on the dataset from DeepMirTar, we achieved a higher accuracy (0. 9787 vs. 0.9348) and F-Score (0.9786 vs. 0.9348) relative to those of DeepMirTar. Similarly, after training on the miRAW training set, we obtained a higher accuracy (0.965 vs. 0.935) and F-Score (0.965 vs. 0.935) relative to miRAW. Finally, we combined the data from DeepMirTar and miRAW, and trained a unified model, named miTAR, which still yields a higher accuracy and F-Score at 0.9545 and 0.9544, respectively.

## Methods and Materials

### Datasets

We collected two datasets that contain sequences of miRNAs and genes from the studies of DeepMirTar and miRAW(Pla, et al., 2018; Wen, et al., 2018). The first dataset was downloaded from the supplementary tables of the DeepMirTar study, which contains 3,963 positive pairs of miRNA:target and 3,905 negative pairs of miRNA:target. The positive pairs in the DeepMirTar dataset were obtained from three resources: mirMark data(Menor, et al., 2014), CLASH data(Helwak, et al., 2013), and PAR-CLIP data(Hafner, et al., 2010). And only the target sites located in 3’UTRs, and the target sites with canonical seeds (exact W–C pairing of 2–7 or 3–8 nts of the miRNA) and non-canonical seeds (pairing at positions 2–7 or 3–8, allowing G-U pairs and up to one bulged or mis-matched nucleotide) were included. The negative pairs were generated by shuffling the real mature miRNAs. Details on generating the datasets are in the DeepMirTar study(Wen, et al., 2018). We further removed the miRNAs that cannot be found from the current version of miRBase (release 22) {http://mirbase.org/ftp.shtml}. Finally, a total of 3,956 positive pairs and 3,898 negative pairs were kept. In the DeepMirTar study, the positive pairs from PAR-CLIP data (48 pairs) were used as an independent test dataset. In our study, we also kept the same 48 positive pairs and further randomly selected an additional 48 negative pairs as an independent test dataset. This set was termed DeepMirTarIn. The remaining data were termed DeepMirTar (Table 1).

**Table 1.** The number of miRNA:target in each dataset used in training, validating and testing the proposed approach.

| Datasets | Positive Pairs | Negative Pairs | Training Set | Validation Set | Test Set |
| --- | --- | --- | --- | --- | --- |
| DeepMirTar dataset | 3,908 | 3,850 | 4,964 | 1,241 | 1,552 |
| miRAW dataset | 31,660 | 30,993 | 40,096 | 10,025 | 12,531 |
| MirTarRAW dataset<sup>a</sup> | 13,860 (3,465 DeepMirTar + 10,395 miRAW) | 13,860 (3,465 DeepMirTar + 10,395 miRAW) | 17,740 | 4,435 | 5,544 |
| DeepMirTarIn dataset<sup>b</sup> | 48 | 48 |  |  |  |
| miRAWIn dataset<sup>b</sup> | 1,000 | 1,000 |  |  |  |
| DeepMirTarLeft dataset<sup>b</sup> | 443 | 385 |  |  |  |
| miRAWLeft dataset<sup>b</sup> | 21,265 | 20,598 |  |  |  |

a: Dataset generated by combining the DeepMirTar and the miRAW dataset. Details are in **Methods and Materials** section.
b: Independent test dataset for validating the method.

In the miRAW study, Albert Pla *et al*.(Pla, et al., 2018) collected a large amount of verified data that included both canonical and non-canonical miRNA:target pairs. The experimentally validated positive and negative miRNA:target pairs were collected from two resources: Diana TarBase(Vlachos, et al., 2015) and MirTarBase(Chou, et al., 2016), and the target site sequences were obtained by cross-referencing with PAR-Clip(Grosswendt, et al., 2014), CLASH(Helwak, et al., 2013), and TargetScanHuman 7.1(Agarwal, et al., 2015). In total, 33,142 positive and 32,284 negative pairs were collected. We further removed the miRNAs that cannot be found from the current version of miRBase. Finally, a total of 32,660 positive and 31,993 negative pairs were retained. In the miRAW study ∼1,700 pairs were used as an independent test dataset. Similarly, we randomly selected 2,000 pairs, including 1,000 positive and 1,000 negative pairs, as an independent test dataset and labelled these miRAWIn. The remaining pairs were labeled miRAW(Table 1).

In addition, we combined ∼33% of miRAW data (20,790 pairs) and ∼90% of DeepMirTar data (6,930 pairs) into one dataset (termed MirTarRAW) for training a unified model. The remaining data from DeepMirTar were taken as an independent test dataset and labelled DeepMirTarLeft; the remaining data from miRAW were taken as an independent test dataset and labelled miRAWLeft (Table1).

For each dataset, we concatenated the sequences of miRNAs from 3’->5’ with their target sequences from 5’->3’. To keep all the miRNA sequences the same length, those miRNA sequences with length less than the longest miRNA (26 nts across both datasets) were padded with ‘N’s. The same padding was done for the target sequences, which had targets sites of up to 53 nts. All the target sites in the miRAW dataset were trimmed to the same length by Albert Pla *et al.*(Pla, et al., 2018), which is 40 nts. After padding, the miRNA sequence and the target sequence were concatenated directly. Thus, the length of the sequences after concatenation for the DeepMirTar dataset is 79, and 66 for the miRAW dataset.

### Overview of our hybrid deep learning-based approach for predicting miRNA targets

Six layers were used for miRNA target prediction (Figure 1). The first layer is an embedding layer. The embedding layer converts the input data into a five-dimensional dense vector that can be initialized randomly and trained with the other five layers. The second layer is a 1D-convolutional layer, which aims to learn the spatial features between miRNA:target. The third layer is a max pooling layer that normally follows the CNN layer to reduce the dimensionality of the input data. The fourth layer is a bi-directional RNN (BiRNN). The BiRNN can learn the sequential features of miRNA:target from the forward and reverse directions. The fifth and sixth layer are dense layers that were used to calculate the final classification. To reduce the probability of overfitting and to make the approach more generalized for predicting future cases, a dropout was added following the second layer (Figure 1). The major elements of the approach are described below. Implementation of the approach was done in Python v3.6.5 using Keras with TensorFlow as the backend. Source codes are available at https://github.com/tjgu/miTAR.

**Figure 1.**
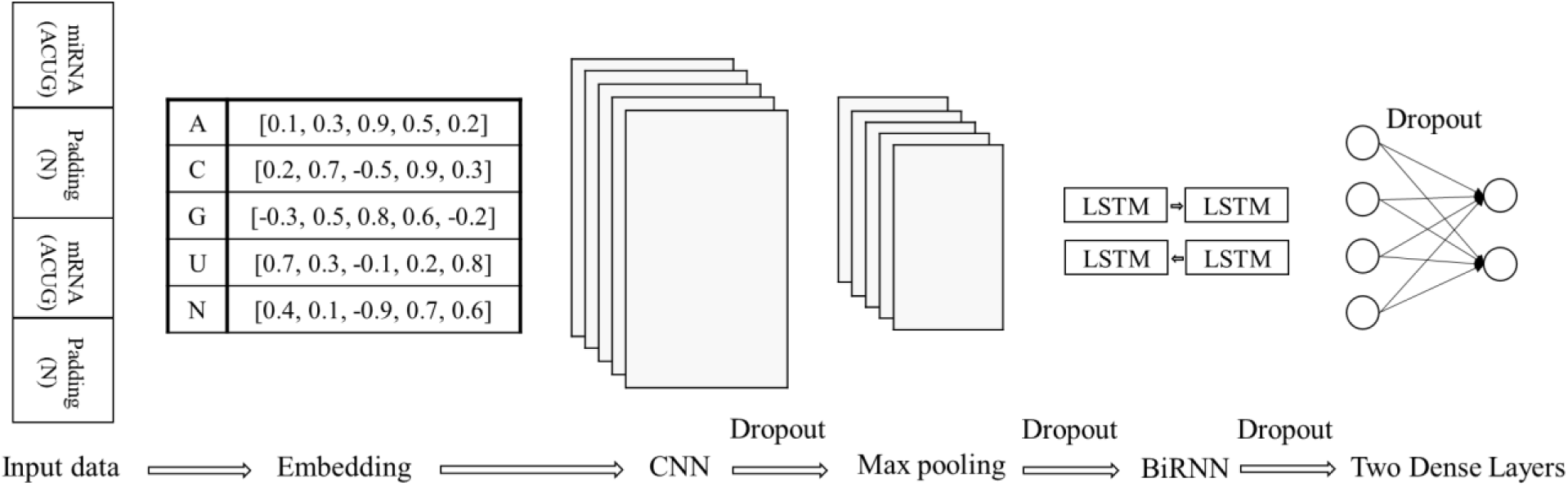
Overview of the proposed deep learning-based approach for predicting miRNA targets. As inputs of the approach, miRNA sequences and gene sequences are padded separately and then concatenated directly. The approach contains one embedding, CNN, Max pooling, and BiRNN layer, and two dense layers. To prevent overfitting, a dropout is added following CNN, Max pooling, BiRNN and the first Dense layer.

### The embedding layer

The one-hot encoding technique is widely used to transform the input sequences into numeric vectors. However, one-hot encoding normally generates sparse high-dimensional vectors that may affect the performance of the model(Taehoon Lee, 2015). The embedding layer can not only transform the sequences into dense vectors, but also can be updated along with all the other layers throughout the training process. It is reported that the embedding layer performs better than one-hot encoding(Hill, et al., 2018). Normally the size of the vector equals the vocabulary size. Since five different letters, {A, U, G, C, N}, were in our datasets, we transformed our input sequences into five-dimensional vectors with one vector for one letter in one input sequence. An example was shown in Figure 1, for instance, for a sequence with length of ten, ten dense vectors are generated and the length for each vector is five.

### The CNN layer

CNN is a neural network that uses filters/kernels to scan the input data in order to capture the embedding spatial information(Zhang, et al., 2019; Zou, et al., 2019). It has been widely used in image processing and recently also been applied in many biological and clinical data analyses. The parameters in a filter/kernel can be shared while scanning different regions of input data. Thus, the model parameters can be greatly reduced, which is one of the advantages to use CNN. In our model, a CNN layer was added following the first embedding layer. The number of kernels was set at 320 with the kernel size of 12. The nonlinear activation function, rectified linear unit (ReLU), was chosen in the CNN layer, which is more robust to gradient vanishing or gradient explosion.

### The RNN layer

RNN is a type of neural networks used widely in natural language processing, speech and image recognition (Zhang, et al., 2019; Zou, et al., 2019). In recent years, RNN has been applied in various biological fields (REF?). The design of RNN naturally fits sequential or time-series data and can model sequences of various length. The hidden layers of an RNN accept not only the input data from previous layers but also the output from the latest time point. A simple RNN can be expanded along the time series into a complicated network. Consequently, a simple RNN is prone to problems like gradient vanishing in the training process and it can be difficult to learn long term dependencies. A few advanced RNNs including long short-term memory (LSTM) and gated recurrent unit (GRU) have been developed to solve these problems. Both approaches use memory based hidden units rather than simple perceptron hidden units, which greatly improve the performance. In our approach, we used the LSTM layer to learn the dependencies between miRNA:target. Since it is possible that the dependencies may exist in the order of target:miRNA, we used bidirectional LSTM (BiLSTM) to learn the sequential information from both directions. The size for the hidden units were set at 32. We also used the ReLU as the activation function for the BiLSTM.

### Dropout

Overfitting is a major problem that deep learning methods face. Dropout is one method to prevent overfitting (Zhang, et al., 2019) and was used in our model. It discards some neuron units from the network according to a certain probability. A variety of probabilities were tested in model selection process. We applied a dropout following every layer, except the first embedding and the last layer.

### Model hyperparameter optimization

To obtain the optimal model parameters, we tested multiple parameters at wide ranges: the learning rates at 0.2, 0.1, 0.05, 0.01, 0.005, and 0.001; the dropout rates at 0.1, 0.2, 0.3, 0.4, and 0.5; and the batch sizes at 10, 30, 50, 100, and 200. The size of epoch was set at 1000. To prevent overfitting, we employed early stopping in addition to dropout. The program stops training when the accuracy of the model does not improve by 0.1% in 100 epochs.

## Results

### Performance comparison with the latest studies using test datasets

We first trained our model on the two datasets separately, DeepMirTar and miRAW. To obtain the optimal model parameters, a wide range for each parameter was tested as described in **Methods and Materials**. We split DeepMirTar and miRAW datasets into three sets separately: 20% were used as a test dataset, 64% were used as a training dataset and 16% were used for a validation dataset (Table 1). For the DeepMirTar dataset the parameters that generated the highest accuracy are learning rate at 0.005, dropout at 0.2 and batch size at 30, which were chosen in the downstream analysis. The model trained with this set of parameters was labelled as miTAR1. Then we randomly split the DeepMirTar dataset 30 times into training, validation and test sets and ran the same model structure 30 times. We obtained an average accuracy of 97.87%, which is higher than the accuracy of 93.48% reported in the DeepMirTar study(Wen, et al., 2018) (Table 2). The set of parameters that produced the highest accuracy for the miRAW dataset is learning rate at 0.1, dropout rate at 0.4 and batch size at 200. We randomly split the miRAW datasets 30 times similarly as the DeepMirTar dataset and obtained an average accuracy of 96.49%, which is higher than the accuracy of 93.5% reported in the miRAW study(Pla, et al., 2018) (Table 2). We labelled this model as miTAR2.

**Table 2.** Performance evaluation metrics for miTAR1 and miTAR2.

| **Model: dataset** | **Accuracy** | **Sensitivity** | **Specificity** | **F-Score** | **PPV<sup>c</sup>** | **NPV<sup>c</sup>** | **Brier Score** |
| --- | --- | --- | --- | --- | --- | --- | --- |
| DeepMirTar:<br>DeepMirTarRaw<sup>a</sup> | 0.9348 | NA | NA | 0.9348 | NA | NA | NA |
| miTAR1:<br>DeepMirTar Test set | 0.9781 | 0.9648 | 0.9921 | 0.9783 | 0.9922 | 0.9641 | 0.0214 |
| miTAR1:<br>DeepMirTar (30 times)<sup>b</sup> | 0.9787<br>[0.9774-0.9801] | 0.9717<br>[0.9694-0.9739] | 0.9857<br>[0.9841-0.9876] | 0.9786<br>[0.9773-0.9800] | 0.9858<br>[0.9840-0.9875] | 0.9719<br>[0.9694-0.9743] | 0.0193<br>[0.0181-0.0206] |
| miRAW:<br>miRAWRaw<sup>a</sup> | 0.935 | 0.935 | 0.938 | 0.935 | NA | NA | NA |
| miTAR2: miRAW Test set | 0.9654 | 0.9609 | 0.9697 | 0.9652 | 0.9695 | 0.9613 | 0.0283 |
| miTAR2: miRAW (30 times)<sup>b</sup> | 0.9649<br>[0.9641-0.9656] | 0.9616<br>[0.9603-0.9629] | 0.9683<br>[0.9670-0.9695] | 0.9651<br>[0.9644-0.9659] | 0.9687<br>[0.9675-0.9700] | 0.9610<br>[0.9597-0.9623] | 0.0271<br>[0.0266-0.0276] |

NA means the value not reported in the corresponding study.
a: DeepMirTarRaw and miRAWRaw present the dataset used in the DeepMirTar and the miRAW study. The best performance values are selected for the performance metrics if multiple values are reported in the respective study for different conditions.
b: Evaluation was done by randomly running on the DeepMirTar and the miRAW datasets 30 times. The average value and the 95% confidence interval (given in []) were reported here. Details are in Results section.
c: PPV represents, positive predictive value; NPV presents negative predictive value.

In addition to the accuracy, we calculated other metrics to evaluate our models’ performance, including sensitivity, specificity, F-measure (or F-score), positive predictive value (PPV), negative predictive value (NPV), and Brier Score. Sensitivity and specificity measure the power of the model for detecting true positives and true negatives, respectively. PPV and NPV measure the precision of the model on correctly predicting the true positives and true negatives, respectively. F-score is the harmonic mean of PPV (precision) and sensitivity (recall), which can be used as a parameter that integrates precision and recall and is more informative for measuring the performance of models. Brier Score measures the accuracy of probabilistic predictions. Rather than using the discrete prediction outcomes, Brier Score uses the prediction probability that is assigned to each outcome. It calculates the mean squared difference between the prediction probabilities and the true values. The results for each model discussed above were shown in Table 2. The sensitivity, specificity, F-score, PPV, and NPV were all higher for our models than the models reported in the DeepMirTar and miRAW studies.

### Performance comparison with the latest studies using independent datasets

Furthermore, we compared the results using the independent test datasets with the studies of DeepMirTar and miRAW. We identified 36 of 48 positive miRNA:target pairs, which is more than the 24 reported in the DeepMirTar study (Table 3). We obtained an accuracy of 96.95 on the dataset of miRAWIn, which is higher than the accuracy of 91.3 reported in miRAW (Table 4).

**Table 3.** Performance comparison between miTAR1 and DeepMirTar on an independent dataset. NA means the value not reported in the corresponding study. The number ahead of / is the number of miRNA:targets identified by the respective models; the number following / is the total number of miRNA:targets in the respective independent datasets. The positive pairs for both models are the same set of pairs. No negative pairs are used in the DeepMirTar study.

| **Model: dataset** | **Identified positive**<br>**miRNA:targets /Total number**<br>**of positive pairs** | **Identified negative**<br>**miRNA:targets/Total number of**<br>**negative pairs** |
| --- | --- | --- |
| DeepMirTar:<br>DeepMirTar<br>Independent test set | 24/48 | NA |
| miTAR1: DeepMirTarIn | 36/48 | 47/48 |

**Table 4.** Performance comparison between miTAR2 and miRAW on an independent test dataset.

| Model:dataset | Accuracy | Sensitivity | Specificity | F-Score | PPV<sup>b</sup> | NPV<sup>b</sup> | Brier Score |
| --- | --- | --- | --- | --- | --- | --- | --- |
| miRAW:<br>miRAWRaw<sup>a</sup> | 0.913 | 0.931 | 0.363* | 0.954 | NA | NA | NA |
| miTAR2: miRAWIn | 0.970 | 0.960 | 0.979 | 0.969 | 0.979 | 0.961 | 0.025 |

NA means the value not reported in the corresponding study.
a: miRAWRaw presents the dataset used in the miRAW study. The best performance values were selected from the reports of the miRAW study.
b: PPV represents positive predictive value; NPV represents negative predictive value.

### Performance comparison with earlier studies using test datasets

Beside DeepMirTar and miRAW, we compared our approach to two other published deep learning methods: deepTarget and miRTDL (Cheng, et al., 2016; Lee, et al., 2016). They were developed earlier, hence, the training datasets they used are much smaller. 507 target site-level and 2,891 gene-level miRNA:target pairs from mirMark repository (Menor, et al., 2014) were taken as the positive training dataset in the deepTarget study. mirMark repository is part of the DeepMirTar dataset, subsequently, we compared miTAR1 with the results reported from deepTarget study. Although deepTarget performs better on NPV (0.9719 vs. 0.9845), our model outperformed the deepTarget model on accuracy, sensitivity, specificity, F-score, and PPV, especially the PPV (0.9858 vs. 0.8848) and F-score (0.9786 vs. 0.9105) (Supplementary Table 1), indicating miTAR1 is much more effective on miRNA target prediction than deepTarget. 1,297 positive and 309 negative miRNA:target pairs from TarBase (Vlachos, et al., 2015) database were taken as the training dataset in the miRTDL study, which is a small portion of the miRAW dataset. Accordingly, we compared miTAR2 with miRTDL. miRTDL only reported their results for accuracy, sensitivity and specificity. Although miRTDL outperforms on sensitivity (96.16 vs. 96.44) slightly, our model performs better on accuracy (96.49 vs. 89.98) and specificity (96.16 vs. 88.43) (Supplementary Table 2).

### Performance evaluation of the unified model, miTAR

Since the length of the input sequences from DeepMirTar (79 nts) and miRAW (66 nts) datasets is different, miTAR1 cannot be fit by miRAW dataset, and miTAR2 cannot be fit by DeepMirTar dataset. To build a unified model, we combined the two datasets and trained a new model that can fit both datasets. Due to much more pairs of miRNA:target in the miRAW dataset, we only randomly extracted ∼33% of miRAW data, which contains 10,395 positive and negative data respectively, but we randomly extracted ∼90% of DeepMirTar dataset, which contains 3,465 positive and negative data separately. We labelled this dataset as MirTarRAW (Table 1). The MirTarRAW dataset were then split into three sets: 20% for testing, 64% for training, and 16% for validation. The remaining data from the DeepMirTar were labelled as DeepMirTarLeft; and the remaining data from the miRAW were labelled as miRAWLeft, which were used as independent test data that never used in the training, validation, or test in the process of selecting the best model parameters. The best model parameters for the combined dataset, MirTarRAW, are learning rate at 0.005, dropout rate at 0.2 and batch size at 100. We randomly split the dataset 30 times and obtained an average accuracy of 95.45%, which is much higher than the accuracies reported in the DeepMirTar (93.5%) study and the miRAW (93.48%) study. We also tested the model using two independent datasets: DeepMirTarLeft and miRAWLeft. The accuracies obtained from both independent datasets are also higher than the reports from either DeepMirTar or miRAW (Table 5). Consistent better results were obtained for all the other metrics tested in our study, including sensitivity, specificity and F-Score (Table 5). These results suggest that our new model is a more accurate model than the current ones. We labelled this model as miTAR and supplied the source code and executable at https://github.com/tjgu/miTAR.

**Table 5.**
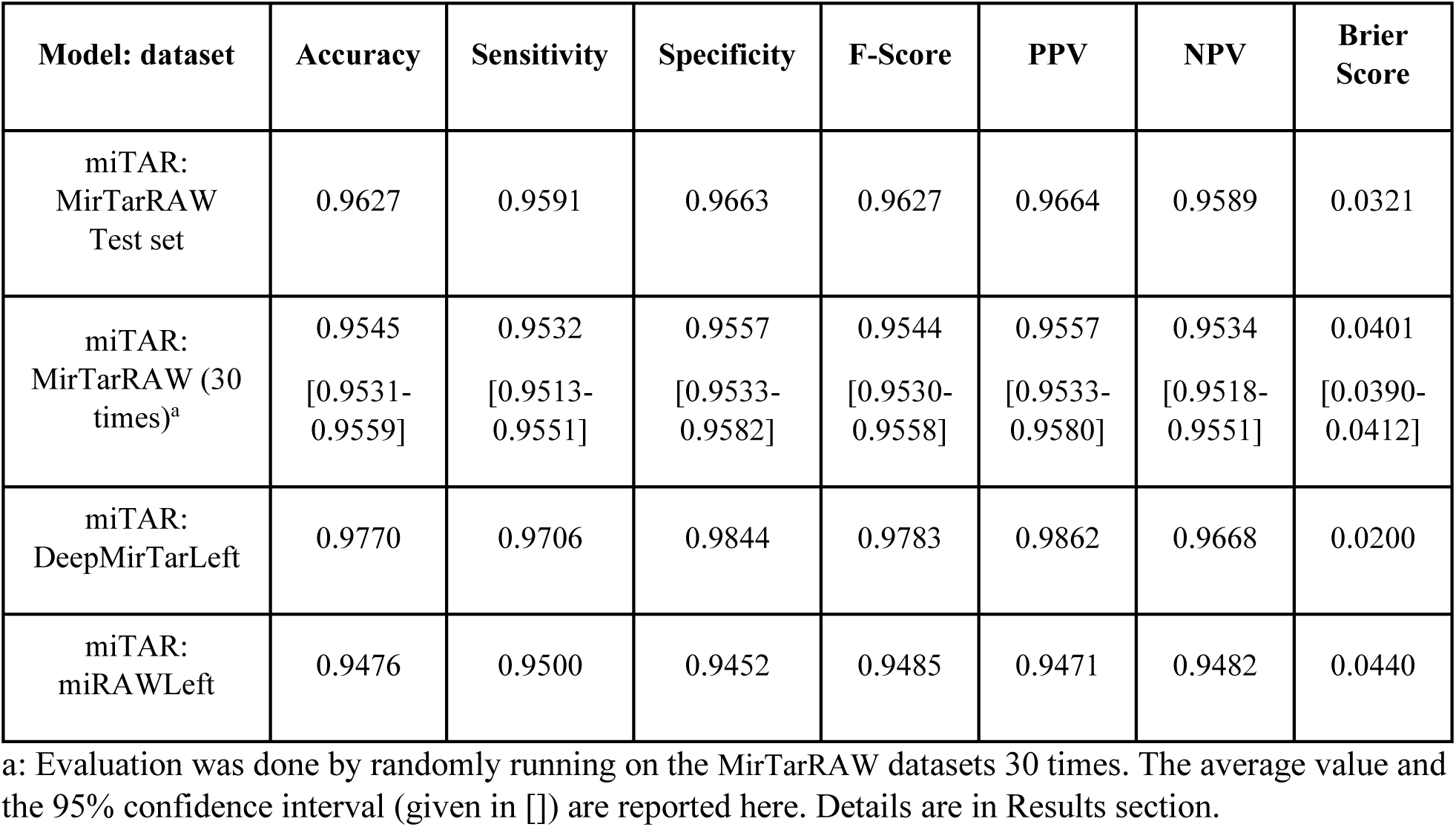
Performance evaluation metrics for the unified model, miTAR using MirTarRAW test datasets and two independent test sets.

## Discussion

We developed a hybrid deep learning-based approach that integrated two major type of neural networks, CNN and RNN, to predict the targets of miRNAs at a higher accuracy than previous methods/tools. Using multiple model evaluation metrics, we demonstrated that our hybrid method significantly outperforms the latest deep learning methods, DeepMirTar and miRAW. Another advantage of our method is that the inputs of our models do not require pre-defined features, which can save users from calculating a large number of features.

The training datasets are more important than we anticipated. We observed a better performance for all models that trained on the DeepMirTar dataset than on the miRAW dataset. For example, when training the model on the two datasets independently, the accuracy for the DeepMirTar dataset is 97.87%, while the accuracy for the miRAW dataset is 96.49% (Table 2). After combining ∼2 fold more data from the miRAW dataset than the DeepMirTar dataset to train the miTAR model (20,790 pairs of miRNA:target vs. 6,930) we still obtained a higher accuracy for the testing data from DeepMirTar (DeepMirTarLeft: 0.9770) than the testing data from miRAW (miRAWLeft: 0.9476) (Table 5). Several factors may potentially impact the performance of the models on the two datasets: the non-canonical miRNA:target pairs; the approaches for generating the negative datasets; the length of the target sequence; noises in the datasets. The miRAW dataset included all possible validated pairs of miRNA:target. Although DeepMirTar dataset included both canonical and non-canonical validated miRNA:target pairs, the non-canonical miRNA:target were limited to the non-canonical seed target sites, which may be easier to learn. Second, the negative pairs from DeepMirTar were generated by shuffling the real miRNA sequences; while the negative pairs from miRAW were generated using the sequences from the genes that were experimentally validated to not be miRNA targets. These differences may add variability to the performance. Third, the length of the target sites is apparently different between the two datasets: the median length of the target sites in DeepMirTar is 22; while the length for all target sites in miRAW is 40. The short-extended length of the target sites from miRAW may not provide much benefit. Fourth, both datasets claimed to be experimental validated target sites, but potential noises still exist. For example, some sites were only validated by one approach or only validated by high-throughput methods (the CLASH data that used in both datasets). In addition, miRAW also included target sites from the broadly conserved human sites that were predicted by TargetScan as positive pairs, although the pairs of miRNA:target reported validated by experiments. The prediction from TargetScan may not be as accurate as high-throughput experimental approaches. The noise arising from different sources may contribute differently on the performance. Nevertheless, our analysis results underscore the importance of the training datasets. More efforts will be needed for collecting true positive and negative miRNA:target pairs, and canonical and non-canonical pairs.

Although linear sequences contain spatial information, when miRNAs bind to the genes, they form hybrid secondary structures. Spatial features exist in miRNA:target. The CNN is designed to capture the spatial features. Therefore, theoretically, models with a CNN perform better. We did a simple test to examine this. We removed the steps that associated with CNN, for example, the CNN and max pooling layers. After removing the two layers and the associated dropouts, we observed poorer performance. Specifically, the accuracy dropped from 96.27% to 93.33%, -similar to the accuracies reported in DeepMirTar and miRAW studies. Although performance may increase significantly by changing other parameters, for example, the batch size and the number of units in the remaining layers, our results support the importance of spatial features in miRNA target prediction. One previous study, the miRTDL study, applied CNN in miRNA target prediction, however, the authors only used CNN as a classifier (Cheng, et al., 2016). The miTAR model is the first to use CNN to capture the potential spatial features directly from the sequences of miRNAs and genes.

The accuracy for all our models was higher than 95.5%, which indicates that our models can learn the intrinsic features between miRNA:target. However, understanding the features from the millions of parameters of the model is complicated, especially for our models with input of sequences. Unlike the models using pre-defined features, we cannot alter one of the pre-defined features to test the importance of each feature. Nevertheless, many known features were reported that are important for a miRNA recognizing its target genes. We can correlate the known features with the output of each layer and validate the importance of the known features. In addition, by scrutinizing the sequences of miRNAs and genes from the positive and negative datasets, new features may be revealed. Attempts to decode the information learned from deep learning models are emerging and will be considered in our future work.

